# A Novel Interaction Between Aquaporin 1 and Caspase-3 in Pulmonary Arterial Smooth Muscle Cells

**DOI:** 10.1101/2023.10.05.561144

**Authors:** Shannon Niedermeyer, Xin Yun, Marielena Trujillo, Manuella Ribas Andrade, Todd Kolb, Karthik Suresh, Mahendra Damarla, Larissa A. Shimoda

**Author notes:** Address for Correspondence: Larissa A. Shimoda, PhD, Division of Pulmonary and Critical Care Medicine, Johns Hopkins University, 5501 Hopkins Bayview Circle, 4A.52, Baltimore, MD 21224.

## Abstract

Pulmonary arterial hypertension (PAH) is a disease in which remodeling of the precapillary pulmonary vasculature leads to hyperplasia and hypertrophy of the muscular vascular wall, and the formation of vaso-occlusive lesions. These pathologic changes are predominantly due to abnormal proliferation and migration of pulmonary arterial smooth muscle cells (PASMCs), enhanced cellular functions that have been linked to increases in the cell membrane protein aquaporin-1 (AQP1). However, the mechanisms underlying increased AQP1 abundance have not been fully elucidated. Here we present data that establishes a novel interaction between AQP1 and the proteolytic enzyme caspase-3. *In silico* analysis of the AQP1 protein reveals two caspase-3 cleavage sites on its c-terminal tail, proximal to known ubiquitin sites. Using biotin proximity ligase techniques, we establish that AQP1 and caspase-3 interact in both HEK293A cells and rat PASMCs. Furthermore, we demonstrate that AQP1 levels increase and decrease with enhanced caspase-3 activity and inhibition respectively. Ultimately, further work characterizing this interaction could provide the foundation for novel PAH therapeutics.

## Introduction

Pulmonary hypertension (PH) is a disease in which remodeling of the precapillary pulmonary vasculature leads to hyperplasia and hypertrophy of the muscular vascular wall, and the formation of vaso-occlusive lesions.^1,2^ These pathologic changes are due in part to abnormal proliferation and migration of pulmonary vascular cells, including pulmonary arterial smooth muscle cells (PASMCs).^3,4^ Ultimately, this remodeling leads to elevated pulmonary vascular pressure and if left untreated, right ventricular (RV) failure and death. Several classes of therapies are available to treat PH, all of which primarily target pathways mediating vasodilation.^5,6^ However, there are no current therapies that target the underlying pathobiology of disease. Efforts to understand the underlying molecular mechanism(s) that drive disease development may provide novel pathway(s) for new therapeutic agents with the potential treat and even reverse disease.

Mounting evidence within the field implicates the protein aquaporin 1 (AQP1) in the development of PH.^7,8^ AQP1 is a 29 kDa transmembrane protein initially described as a water transport channel.^9,10^ Our lab showed AQP1 protein is present throughout the pulmonary vasculature and increased in PASMCs from animal models of PH^11^, a finding confirmed by other labs.^12^ We demonstrated AQP1 is necessary for enhanced proliferation and migration of PH PASMCs, independent of its water channel function.^3,4,11^ Subsequent studies found PH was attenuated in mice with global deletion of AQP1^8^ or when AQP1 was reduced in the lung using silencing approaches.^13^ In rodent models of severe PH, there appears to be an increase in protein abundance without a concomitant increase in mRNA^4^, suggesting either enhanced protein translation or reduced degradation in diseased cells.

The primary degradation pathway for AQP1 is via ubiquination.^14^ Inspection of the primary AQP1 protein structure demonstrated two confirmed ubiquitination sites (K243 and K267) located outside the water transport channel on the small 37 amino acid c-terminal cytosolic tail^15^. Using *in silico* analysis, we also found three potential caspase-3 (casp3) cleavage or interaction sites, denoted by the conformation DXXD (Fig 1A). One of these sites is located on an extracellular portion of the water transport pore (Asp129). The other two potential cleavage sites are within the c-terminal cytosolic tail (Asp236, Asp256), each located proximally to a ubiquitination site (Fig 1B). Identification of potential cleavage sites suggest the possibility of an interaction between AQP1 and casp3.

**Fig 1:**
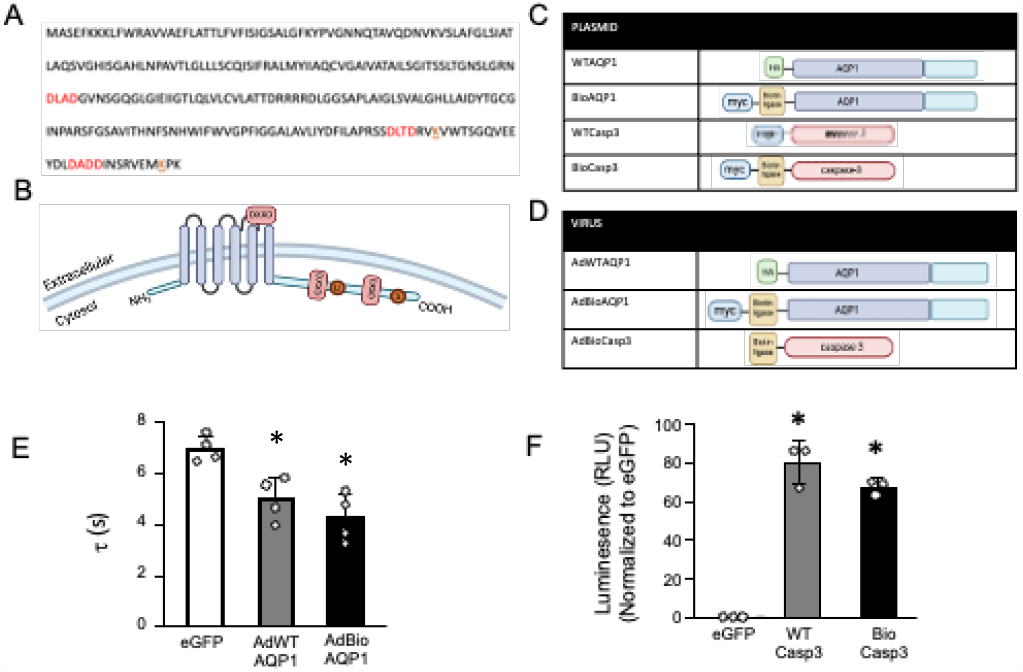
**A)** Primary sequence and **B)** Schematic representation of human aquaporin 1 (AQP1) protein. Potential caspase-3 (casp3) cleavage sites are Indicated In red and ublqultln sites are indicated in orange. Tables showing relevant **C)** plasmid and **D)** adenoviral constructs used in this study. **E)** Bar and scatter graph shows mean.± SO values for the time constant of decay (τ) of calceln fluorescence (quenching) measured In adherent calceln-loaded PASMCs. τ, a measure of water transport, was calculated when perfusate was switched from 140 to 380 mOsm in cells infected with AdGFP (GFP), AdWTAQP1 and Ad8ioAQP1. F) Bar and scatter graph shows mean±SD of caspase-3 activity In HEK293 cells transfeeted with enhanced green fluorescent protein (eGFP, control) or with wild-type (WTCasp3) or biotin llgase-fused (BioCasp3) caspase 3. * indicates significant difference (p<0.001) from eGFP (control) by ANOVA and Holm-Sldak post-hoc test. There was no significant difference between WTCasp3 and BioCasp3 (p=0.139).

Casp3 is a member of the cysteine-dependent aspartate-specific proteases (caspases).^16^ It exists in the cytosol as an inactive dimer.^17,18^ Casp3 is first activated by cleavage of the interdomain linker between the large (17-19 kDa) and small subunits (12 kDa), while a second cleavage of the N-terminal prodomain renders the enzyme fully active. In addition to its highly regulated role as part of the pro-apoptotic caspase cascade, active cytosolic casp3 interacts with other intracellular signaling pathways in non-apoptotic functions. ^16,19–21^

Given the location of the casp3 cleavage sites on the AQP1 c-terminal tail, we hypothesized casp3 interacts with AQP1 and regulates its protein abundance in PASMCs. The present study used proximity biotinylation approaches to assess the interaction between AQP1 and casp3 in both HEK293A cells and PASMCs, as well as the potential implications for AQP1 protein expression.

## Methods

### Ethical Approval

All procedures and protocols in this study were conducted in accordance with NIH guidelines for the proper care and use of animals in research and were approved by the Johns Hopkins University Animal Care and Use Committee.

### Cell Culture

HEK239A cells (Thermo Fisher) were grown to 80-100% confluence in growth media (DMEM 1x with 45g/L D-glucose with 1% penicillin streptomycin, 10% fetal bovine serum, L-glutamine [5ml/1L], and MEM NEAA [5ml/1L]). Rat PASMCs were isolated by enzymatic digestion from distal pulmonary arteries (200-600 μm) and cultured in Smooth Muscle Cell Medium (Sciencell, Cat 1101) supplemented with smooth muscle cell bullet kit (Lonza). Following isolation, cell purity is confirmed by spindle morphology and staining for smooth muscle cell specific markers. Cells were only used at passage 0-2. We routinely test for mycoplasma contamination.

### Designing plasmid and viral constructs

Several plasmids and adenoviral constructs were created for biotinylation experiments (Fig 1C and D). The BirA gene (NCBI WP_010880335.1) was utilized to add a promiscuous biotin ligase tag to proteins as previously described.^22^ Other tags, including Human influenza hemagglutinin (HA) and myc, were utilized to aid in protein identification via immunoblot (see Supplemental Table 1 for additional details).

### Biotin Proximity Ligase Assay

Cells were transduced with protein constructs alone (control) or fused to biotin ligase. Plasmid constructs were used in human embryonic kidney (HEK293A) cells and adenoviral constructs in PASMCs. After 24 h, cell-permeable biotin (50 µM) was added to the media and cells incubated for an additional 12-16 h. Cells were washed with PBS and lysed (input). Biotinylated proteins were precipitated with streptavidin-linked beads. Beads were washed and proteins eluted with sample buffer which includes biotin to facilitate isolation (eluent). The final eluent was immunoblotted for the protein of interest. ^23^

### Immunoblotting

Proteins (5 μg) were separated on 12% SDS acrylamide gels, transferred to polyvinylidene fluoride membranes, and blocked in 5% nonfat dry milk. Membranes were the probed with primary and secondary antibodies and bands visualized by ECL (Amersham) according to the manufacturer’s instructions. Densitometry was used to quantify protein levels. The following specific primary antibodies were used: β-tubulin (loading control; Sigma T7816), aquaporin 1 (Sigma AQP11-A); HA (Cell Signaling 3724S); myc (Cell Signaling 2276S); total caspase 3 (Cell Signaling 9662S); and cleaved caspase 3 (Cell Signaling 9664S). ^11^

### Caspase activity

Activity was measured using a luminescent caspase-3/7 activity assay (Caspase-Glo® 3/7). Cell lysates (5 µg) were combined with proluminescent caspase-3/7 DEVD-aminoluciferin substrate in a 96 well plate, and serial measurements made using a luminometer every 5 min for up to 90 min.

### Water permeability assay

Water permeability was estimated using the calcein self-quenching method.24,25 Primary PASMCs were infected with AQP1-containing adenoviral constructs, trypsinized and re-plated onto glass coverslips. Cells were incubated with calcein (5 µM) for 30 m at 37°C after which coverslips were placed in a closed chamber and perfused with isotonic, hypotonic and hypertonic fluids in rapid succession. Cells were excited with light filtered at 490 nm, and images of emitted fluorescence (515 nM) were captured. Fluorescence was normalized to initial value (F0) and the decrease in calcein fluorescence with the change from hypotonic to hypertonic solution was fit with a single exponential to calculate the time constant (τ τ) for each cell. For all cells within an experiment, τ τ were averaged to obtain a single value per experiment.

### Caspase-3 activation or inhibition

To inhibit casp3, primary cells were pretreated with Z-DEVD-FMK (DEVD, 50 µg/mL) for 1 hr at 37°C. DMSO (0.5%) was used as a vehicle control. After pre-treatment, caspase-3 was activated with hydrogen peroxide (500 µM), with PBS as a control for 24. Cells were collected into cell lysis buffer (80-100 µl of T-PER™, Thermo Fisher and protease inhibitor cocktail) and lysates were immunoblotted for AQP1 as described above.

### Statistical Analysis

Data are expressed as scatter plots with bars representing means ± SD. Each dot represents a separate experimental run. For PASMCs, all experimental runs were performed on cells from different animals; thus, “n” also refers to the number of biologic replicates (animals). All data were tested for normality and equal variance prior to running statistical tests. Data that were not normally distributed were log(10) transformed and retested for normality and equal variance prior to running statistics. Statistical comparisons were performed using Students t-test for data in two groups, or one- or two-way ANOVA with a Holm-Sidak post hoc test for multiple group comparisons.

## Results

### Activity of biotin ligase tagged AQP1 and casp3

PASMCs were infected with AQP1 and casp3 adenoviral constructs containing wild type proteins or proteins with the addition of a biotin ligase to the n-terminus (Fig 1D) to test for proper activity prior to use in experiments. Water transport rate in PASMCs infected with AdWTAQP1 and AdBioAQP1 were similarly increased compared to PASMCs infected with AdGFP (Fig 1E), demonstrating the addition of the biotin ligase does not alter AQP1 water permeability. Similar experiments were performed to assess casp3 activity; however, due to an inability to adequately increase wild-type casp3 in PASMCs (Supplemental Fig 1), casp3/7 activity assays were performed on HEK293A cells transfected with eGFP (control), WTCasp3 and BioCasp3 (Fig 1F). The addition of the biotin ligase tag did not significantly alter caspase activity.

### Biotin Proximity Labeling assays in HEK293A cells

Biotin proximity labeling assays were utilized to evaluate interactions between AQP1 and casp3. HEK293A cells were chosen as the initial expression system given ease of manipulation and stability. First, WTAQP1 and BioAQP1 were expressed in HEK293A cells (Fig 2A), which do not express native AQP1. WTAQP1 was used as a control to evaluate whether the addition of cell permeable biotin alone leads to nonspecific biotinylation of cellular proteins. Immunoblot utilizing the input (cell lysate after transfection and incubation with cell permeable biotin) showed equivalent levels of casp3 in the input lysates, while eluents (cell lysate after incubation overnight with streptavidin beads) exhibited a band for biotinylated casp3 at approximately 32 kDa (the additional weight of biotin being 0.2kD) only in samples containing BioAQP1 (Fig 2B).

**Fig 2:**
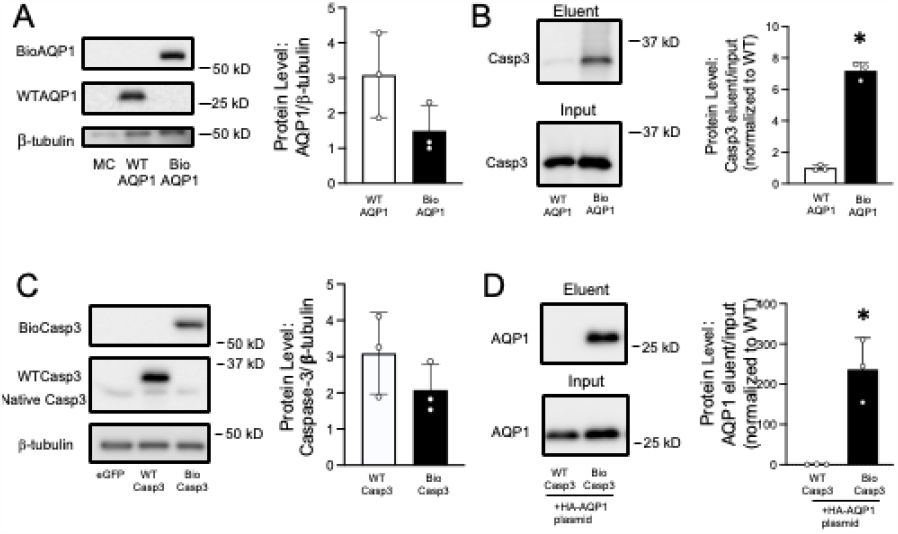
**A)** Representative Image showing expression of wild type (WTAOP1) and biotin llgase conjugated AQP1 (BloAQP1) In HEK293A cells, Which do not express native AQP1. The addition of a biotin ligase increases the molecular weight by ∼35 kD. M-cheny (MC) was used as a control. Bar and scatter graph shows mean±SD values for expressed AQP1 constructs normalized to-1.ubulin (loading control). There was no difference between WTAQP1 and BloAQP1 by two-tailed t-test (p0.121). **B)** Representative Image showing caspase-3 (casp3) protein captured by streplavldln puD down (eluenl) and In total lysates (Input) In HEK293A oells transfected with WTAQP1 and BloAQP1. Bar and scatter graph represents mean±SD ratio values for casp3 In eluenl and Input (loading control). * Indicates significant difference (p<0.001) by two-tailed t-tesl. **C)** Representative Image showing expression of wild type (WTCasp3) and biotin llgase conjugated casp3 (BloCasp3) in HEK293A cells. Enhanced green nuoresoenl protein (eGFP) was used as a control. Bar and scatter graph shows mean±SD of expressed casp3 constructs normalized to β-lubulin. There was no significant dift’erence between groups by two-tailed I-lest (p-0.257). **D)** Representative image showing AQP1 protein captured by streptavidin pull down (eluent) and expression in total lysales (lnput) in HEK293A oeHs transfected with WTCasp3 and BioCasp3. Bar and scatter graph represents mean±SD values for ratio of AQP1 protein in eluent and input. * indicates significant difference (p<0.001) by two-tailed I-lest. For all graphs, individual dots represent experimental replicates.

To confirm bidirectionality, biotinylation experiments were repeated using casp3 constructs (Fig 2C). Once expression was confirmed, biotin proximity labeling was performed using HEK293A cells transfected with wild-type (WTCasp3) or biotin ligase-fused casp3 (BioCasp3). Notably, co-transfection with wild type AQP1 (WTAQP1) was necessary given the lack of endogenous AQP1. In Fig 2D, the input blots demonstrate that AQP1 was expressed in the cells relatively equal amounts. Biotinylation of AQP1 was evident as a strong band in cells transfected with BioCasp3 but not in the cells expressing WTCasp3.

### Biotin Proximity Labeling assays in PASMCs

While an interaction between AQP1 and casp3 was demonstrated in HEK293A cells, we wanted to confirm this interaction in our primary cells of interest. PASMCs were infected with wild type AQP1 (AdWTAQP1, control) or biotin ligase-fused AQP1 (AdBioAQP1) and biotinylation experiments performed. AdWTAQP1 and AdBioAQP1 were equally expressed in PASMCs (Fig 3A). Input lysates for the biotin pulldown showed equal amounts of casp3, while eluents showed only a band for biotinylated casp3 in cells infected with AdBioAQP1 (Fig 3B). Initially eluent was probed with an antibody to total casp3, which would be expected to identify full length (inactive) pro-casp3 (32 kDa) and any smaller cleaved (active) casp3 fragments. With this antibody, only pro-casp3 was identified in the AdBioAQP1 lane. In subsequent experiments, an antibody to cleaved caspase-3 was utilized. After prolonged exposure, faint bands were eventually detected in the AdBioAQP1 lane corresponding to the smaller fragments of cleaved casp3 (Supplemental Fig 3A and 3B).

**Fig 3:**
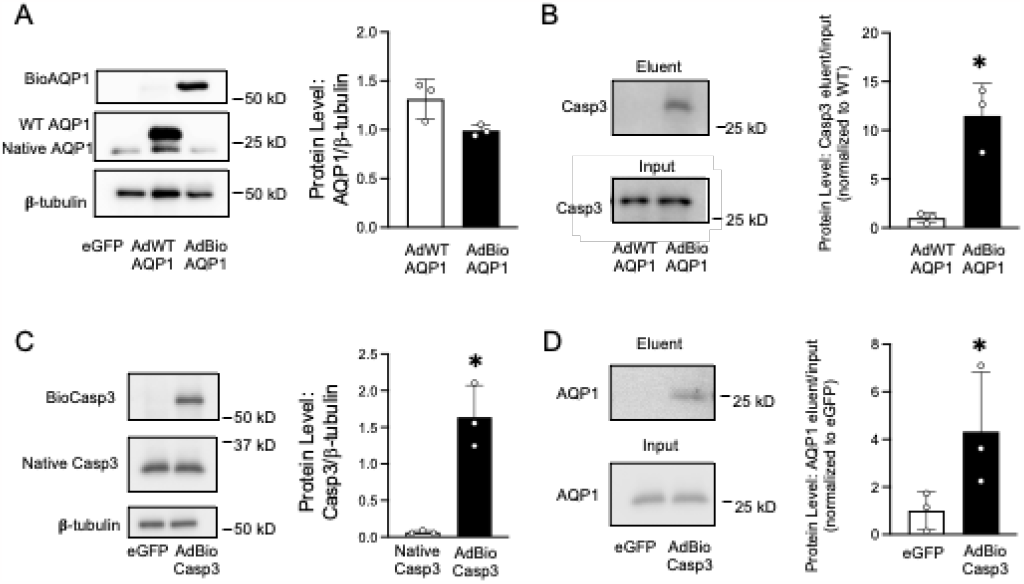
**A)** Representative image showing expressed wild type (WTAQP1) and AQP1 fused to biotin ligase (BioAQPI) In PASMCs (probed with HA). Samples are from the same blot but cropped for size. Bar and scatter graph shows mean±SD values for expressed AQP1 constructs normalized to p-tubulin (loading control). There was no significant difference between AdWTAQPI and AdBioAQPI by two-tailed t-test (p=0.098). **B)** Representative image showing caspase-3 (casp3) protein captured by streptavidin pull down (eluent) and in total lysates (input) in PASMCs infected with WTAQP1 and BioAQPI. Bar and scatter graph represents mean±SD of casp3 in eluent. * indicates significant difference (p=0.006) by twotailed t-test. **C)** Representative image showing expression of biotin ligase-conjugated casp3 (BioCasp3) in PASMCs, with eGFP used as a control. Bar and scatter graph shows mean+SD of casp3 protein. * indicates significant difference (p=0.003) by two-tailed t-test **D)** Representative image showing AQP1 protein captured by streptavidin pull down (eluent) and in total lysates (input) in PASMCs infected with Bk>Casp3. Bar and scatter graph represents mean ± SD of AQP1 in eluent. * indicates significant difference (p=0.047) by one-tailed t-test. For al experiments, individual dots represent expenmental replicates in cells isolated from different animals.

In complementary experiments, adenoviral casp3 constructs were utilized in biotinylation experiments. Figure 3D is a representative blot of AdBioCasp3 experiments probed for AQP1. Surprisingly, while bands for BioCasp3 were clearly visible (Fig 3D), we had difficulty finding expression of WTCasp3 in our cells (Supplemental Fig 1), despite significant amounts of WTCasp3 being expressed in HEK239 cells using the same construct in plasmid form. Total amount of casp3 in the input was similar between cells infected with AdBioCasp3 and cells infected with a control adenovirus (AdeGFP); however, only the eluent of cells infected with AdBiocasp3 showed biotinylation of AQP1 (Fig 3E).

### Effect of Caspase-3 activation and inhibition on AQP1 protein

Given that potential casp3 cleavage sites were identified upstream of lysine residues corresponding to ubiquitination sites, we speculated that interaction with casp3 might increase AQP1 abundance. To evaluate the hypothesized functional relationship between AQP1 and casp3, control PASMCs were pretreated for 30 min with either DEVD (50 µg/mL) or DMSO (vehicle) and then exposed to an oxidant, hydrogen peroxide (H_2_O_2_; 500 µM), for 24 h to activate casp3. Cells treated with PBS were used as a control. Initial experiments showed caspase3/7 activity was increased in response to H_2_O_2_ (Fig 4A), a response that was completely blocked by pretreatment with DEVD. Next, we measured AQP1 protein expression. In vehicle-treated cells challenged with H_2_O_2_, AQP1 protein levels were higher than in PBS-treated controls. In cells pre-treated with DEVD to block casp3 activity, AQP1 protein levels were unchanged by H_2_O_2_ (Fig 4B and C).

**Fig 4:**
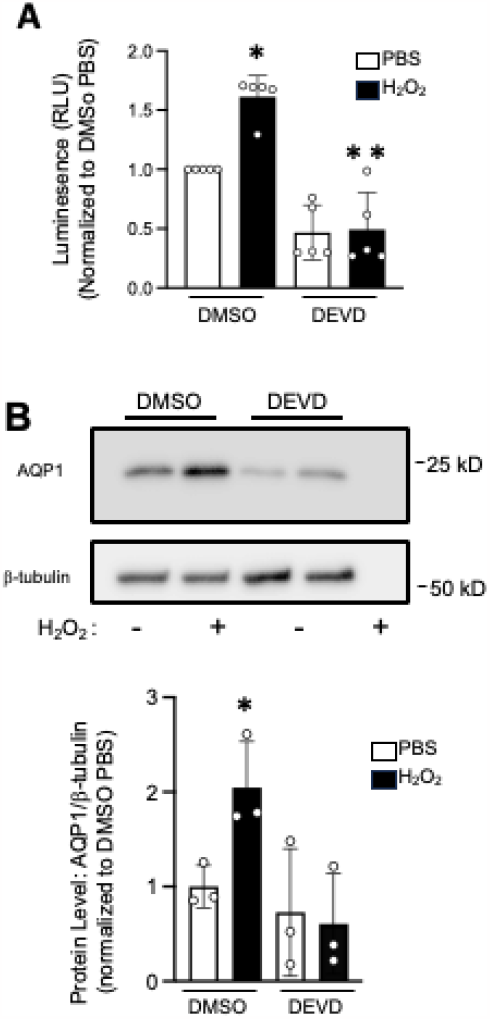
**A)** Bar and scatter graph shows mean±SD caspase-3/7 activity in PASMCs treated with the caspase-3 (casp3) inhibitor DEVD (50 pg/mL) or DMSO (vehicle) for 1 h and then challenged with hydrogen peroxide (H2O2; 500 pM; 24 h) or PBS (vehicle). * indicates significant difference from DMSO PBS (p<0.012) and * * indicates significant difference (p=0.002) from DMSO H_2_O_2_ by two-way ANOVA with Holm-Sidak post-hoc test **B)** Representative image showing AQP1 and β-tubulin (loading control) protein in cells pre-treated with DEVD or DMSO and then challenged with H_2_O_2_ or PBS. Bar and scatter graph shows mean± SD of AQP1 protein.* indicates significant difference (p<0.05) by two-way ANOVA. For all graphs, individual dots represent experimental replicates in cells isolated from different animals.

## Discussion

In this report, using bi-directional biotin proximity ligase assays, we demonstrated a novel interaction between AQP1 and casp3 in both a HEK293A cell line and in primary PASMCs. We also show activation of casp3 enhanced AQP1 protein levels, a response that could be blocked by inhibiting casp3 activity, suggesting a functional role for this interaction in regulating AQP1 abundance.

The mechanism by which AQP1 protein levels are regulated remains poorly understood. Spurred by questions as to how AQP1 abundance is controlled in PASMCs in the absence of a change in mRNA^4^, we explored the AQP1 protein sequence to identify potential ubiquitin sites. Use of publicly available databases (UniProt and PhosphoSitePlus) revealed ubiquitination sites in human AQP1 at residues K243 and K267 in the cytosolic cterminal region of the protein.^15^ *In silico* analysis revealed the presence of residues corresponding to the known casp3 recognition sequence (DXXD) found in many casp3 substrates directly upstream of both ubiquitination sites.^16^ The presence of such motifs suggested the possibility that casp3 may interact with and/or cleave AQP1. We confirmed interaction using proximity biotin ligase assays in both HEK293A cells and PASMCs.

Interestingly, based on the size of the bands visualized for biotinylated capase-3 (Fig 2B and Fig 3B), the predominant form of casp3 interacting with AQP1 appears to be (inactive) pro-casp3. This is consistent with results from complimentary experiments in which casp3 constructs were used, all designed with the biotin ligase fused to the N-terminal of casp3, proximal to the prodomain, which is cleaved upon casp3 activation.^18^ If only active casp3 interacted with AQP1, no biotinylation of AQP1 would have been detected. This finding does not rule out the possibility that active casp3 interacts with AQP1, as identification of small casp3 fragments can be challenging in these assays secondary to antibody specificity and large amount of protein required to detect smaller fragments. Upon further probing with an antibody specific for cleaved (active) casp3 and using prolonged exposure times, we were able to detect faint bands correlating with the size of cleaved casp3 fragments (Supplemental Figure 3). Thus, while both forms of casp3 appear to interact with AQP1, the predominating form is pro-casp3, potentially a reflection of the relative amounts of pro-casp3 (high) and active cleaved casp3 (low) within cultured PASMCs under basal conditions.

Mounting evidence suggests that sub-apoptotic levels of cleaved (active) casp3 may play a role in a variety of cellular functions via cleavage of pathway substrates.^19,26^ There is not however, to our knowledge, a biologically necessary threshold amount of casp3 activity required to participate in these non-apoptotic pathways. Thus, it is plausible that even the very low levels of active casp3 detected in our biotinylation experiments participate in regulating cell function. We hypothesized, given the location of the casp3 cleavage sites on the AQP1 cytosolic tail, that cleavage of tail fragments may increase AQP1 protein abundance, likely due to loss of ubiquitination. Indeed, inhibiting casp3 activity with DEVD at baseline reduced the amount of AQP1 protein in 2 of 3 samples tested. By enhancing casp3 activity with addition of H_2_O_2_, we were able to clearly identify changes in AQP1 protein levels within 24 h, a response that was completely prevented by inhibiting casp3. Our results, while preliminary, suggest a novel mechanism by which AQP1 may circumvent ubiquitination and degradation, prolonging protein abundance.

Given the preliminary nature of this report, there are some limitations. BioID is an excellent tool for identifying protein-protein interactions, especially those that are transient or weak, but AQP1/casp3 interactions should also be confirmed in future experiments by other methods (FRET, co-immunoprecipitation, etc). Also, while the easy detection of inactive (pro-) casp3 in our biotinylation experiments strongly suggests the interaction site is distinct from the casp3 cleavage sites, we have not yet identified the portion of the protein responsible. Finally, while our results indicate casp3 activation is correlated with AQP1 abundance, we have not yet measured ubiquitination of AQP1 when casp3 is activated or inhibited nor proven that casp3 cleaved AQP1. Thus, we cannot state with certainty that cleavage proximal to the ubiquitination sites is the mechanism by which casp3 increases AQP1 protein. We also have not explored whether interrupting the interaction between casp3 and AQP1 liberates casp3 and increases its activity. These are important and exciting avenues for future exploration.

In summary, this work establishes a novel interaction between AQP1 and casp3, and a new line of inquiry regarding the regulation of AQP1 protein abundance by casp3 in PASMCs. Elevations in AQP1 have been explicitly linked to increased migration and proliferation in PASMCs^4,11,13^ contributing to the pathologic remodeling in PH.^27^ While transcriptional upregulation in PH models likely contributes to increased AQP1 expression *in vivo*, decreased protein degradation may also play an important role. Ultimately, disruption of the AQP1/casp3 interaction may provide a potential novel target for new PH therapies to treat and potentially reverse disease.

This work was supported by NIH grants F32HL165766-01, R01 HL159906 and R01 HL126514 and the Bauernschmidt Fellowship Award.

