## Supplemental Figures for "A Novel Interaction Between Aquaporin 1 and Caspase-3 in Pulmonary Arterial Smooth Muscle Cells"

**Supplementary Data:**

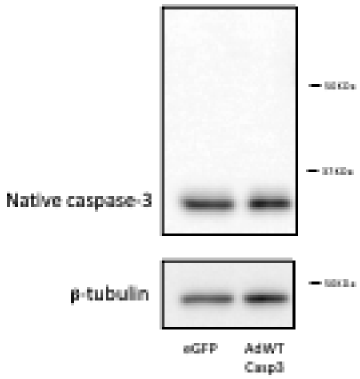


**Fig 1**: Representative blot showing expression of enhanced green fluorescent protein (eGFP, control), wild type caspase-3 (AdWTCasp3). Both viruses infected using 50 infectious units/cell. There is minimal to no expression of AdWTCasp which we would expect to see as a band slightly higher than native caspase-3 .

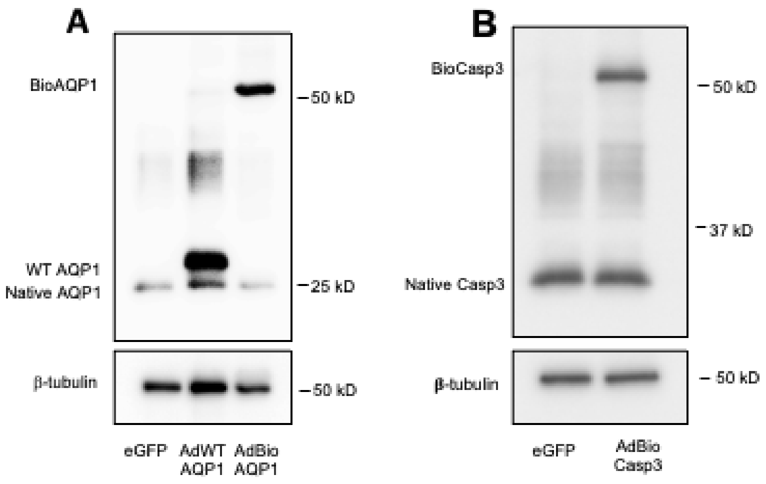


**Fig 2: A)** Representative image showing expressed wild type AQP1 (AdWTAQP1) and AQP1 fused to biotin ligase (AdBioAQP1) in PASMCs, probed for AQP1. **B)** Representative image showing expressed enhanced green fluorescent protein (eGFP) and caspase-3 fused to biotin ligase (AdBioCasp3) in PASMCs, probed for caspase-3.

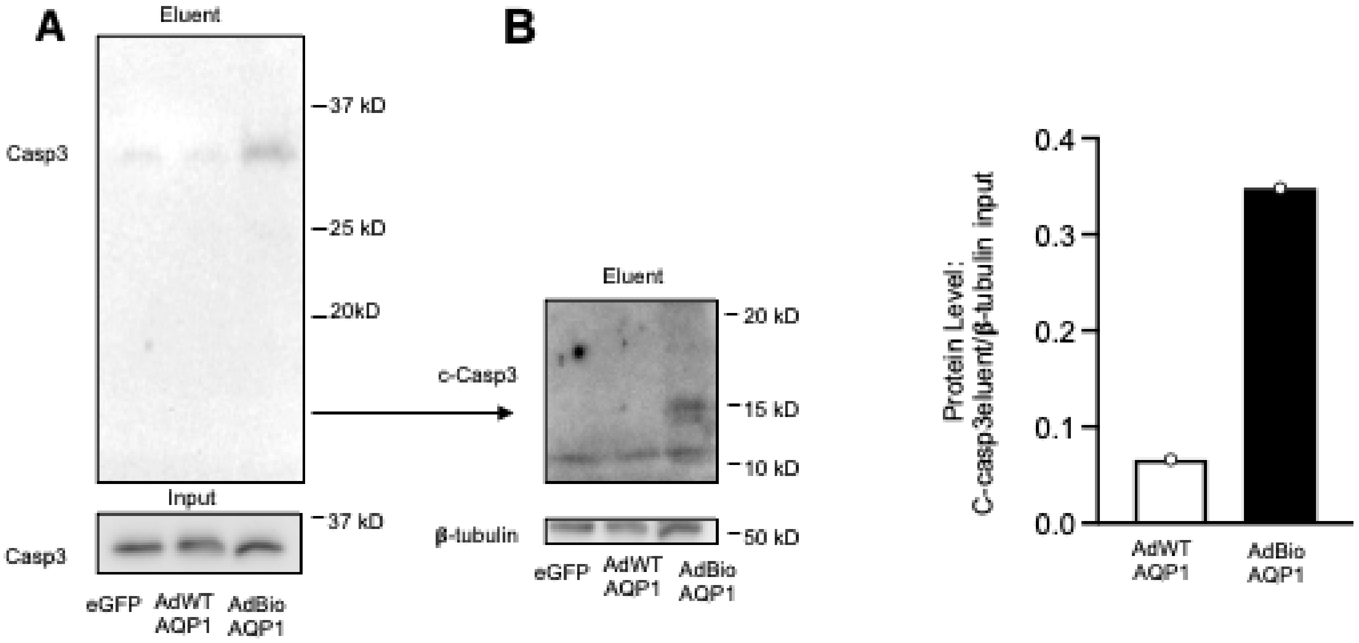


**Fig 3: A)** Representative image showing caspase-3 (casp3) protein captured by streptavidin pull down (eluent) and in total lysates (input) in PASMCs infected with WTAQP1 and BioAQP1. **B)** The same samples showing cleaved caspase-3 (c-casp3) after re-probing with cleaved-caspase3 and prolonged exposure. Bar and scatter graph shows c-casp3 protein as a ratio to the β-tubulin from the input.
